# Functional characterization of a DNA-dependent AAA ATPase in *SimranZ1*, a F cluster mycobacteriophage

**DOI:** 10.1101/2021.03.11.434926

**Authors:** Ritam Das, Urmi Bajpai

## Abstract

Mycobacteriophages are viruses of *Mycobacterium* spp. with promising diagnostic and therapeutic potential. Phage genome exploration and characterization of their proteomes are essential to gain a better understanding of their role in phage biology. So far, about 2014 mycobacteriophages have been genomically defined and 1563 phage protein families (phamilies) are identified. However, the function of only a fraction (about 15%) is known and a majority of ORFs in phage genomes are hypothetical proteins. In this study, from the annotated genome of a F1 cluster mycobacteriophage *SimranZ1*, a putative AAA ATPase (Gp65, Pham 9410) is characterized as a DNA-dependent ATPase. Sequence-based functional annotation predicted Gp65 to belong to the P-loop NTPase superfamily, having AAA_24 and RecA/RadA domains which are known to be involved in ATP-dependent DNA repair/maintenance mechanism. On molecular docking, Gly21 and Ser23 of Gp65 showed specific binding with ATP. Using a microtiter plate assay, ATPase activity of Gp65 was experimentally verified which was found to increase in the presence of dsDNA. Gel electrophoresis under non-denaturing condition showed the oligomeric states of Gp65 and Transmission Electron Microscopy revealed it to exist as a hexamer having a prominent central pore with a diameter of 1.9 nm. In summary, functional characterization of Gp65 as a DNA dependent AAA ATPase indicates its role in DNA repair/maintenance mechanism in mycobacteriophages.

## 1. Introduction

Mycobacteriophages or viruses that infect and lyse *Mycobacterium* spp. were first isolated in 1947 and have been long studied for the importance of mycobacterial diseases (McNerney, 1999). Due to high specificity, mycobacteriophages were principally used to detect and differentiate between various *Mycobacterium* spp by phage typing, which considerably reduce the time taken to diagnose slow growing pathogens such as *Mycobacterium tuberculosis* (Hatfull, 2019). Apart from phage typing, their ability to infect and lyse host bacterium has urged researchers to study mycobacteriophages for their anti-mycobacterial properties. With increasing reports on the emergence of multiple and extremely drug-resistant *Mycobacterium* species, phage therapy for mycobacterial infections is being considered as a viable option. The recently reported successful phage therapy of a drug-resistant *M.abscessus* infection has instilled more confidence in bacteriophages as antimycobacterial agents (Dedrick *et al*., 2019). However, a large number of genes in phage genomes code for hypothetical proteins (Hatfull, 2019) or experimental validation of the predicted function is not done. Hence, functional annotation of genome of bacteriophages is important for understanding their biology and for their application as therapeutics.

According to the Actinobacteriophage database (https://phagesdb.org/), about 11,473 mycobacteriophages have been reported worldwide, which are classified into 27 clusters (and sub-clusters) and 09 singletons (Pope *et al*., 2015). Based on the similarity in amino acid sequence of the phage proteins, there are 1536 ‘phamilies’ in the database (Hatfull *et al*., 2006). Pham 9410 consists of amino acid sequences that code for AAA ATPases.

AAA proteins (ATPases associated with several cellular activities) form a diverse superfamily, involved in ATP dependent remodelling of macromolecules. Proteins belonging to this superfamily have a conserved ATPase module with an αβα domain, which contains the conserved Walker A and Walker B motifs of the P-loop NTPase (Snider *et al*., 2008). Bacteriophage-encoded AAA ATPases are essentially the terminase enzymes that aid in the packaging of dsDNA in the procapsid during viral assembly, using ATP as an energy source (Hilbert *et al*., 2015). However, in 38 mycobacteriophages belonging to F cluster, we found the presence of an additional AAA ATPase protein besides the large and small subunit of terminase. In *SimranZ1* mycobacteriophage (KY385384.1), which belongs to cluster F1 (Bajpai *et al*., 2018), Gp1 and Gp2 are predicted to code for small and large subunit of terminase, respectively and the additional AAA ATPase is coded by Gp65. In this study, we have carried out structural and functional characterization of Gp65 by using *in silico* tools and experimental data.

## 2. Materials and methods

### 2.1 *In silico* analysis of Gp65 of *SimranZ1* mycobacteriophage

#### 2.1.1 Analysis of Gp65 sequence

The nucleotide and amino acid sequence of Gp65 of *SimranZ1* were retrieved from the NCBI database (Taxonomy ID: 1933771). P-BLAST suite (https://blast.ncbi.nlm.nih.gov/Blast.cgi) (Altschul *et al*., 1990), ClustalW tool (https://www.genome.jp/tools-bin/clustalw) (Thompson *et al*., 1994) and Phylogeny.fr (http://www.phylogeny.fr/) (Dereeper *et al*., 2008) were used for homology studies, multiple sequence alignment and phylogenetic relationships, respectively. Conserved domain analysis was done using NCBI Conserved Domain Search tool (https://www.ncbi.nlm.nih.gov/Structure/cdd/wrpsb.cgi) (Marchler-Bauer *et al*., 2010) and InterProScan 5 (https://www.ebi.ac.uk/) (Madeira *et al*., 2019).

#### 2.1.2 Characterization of Gp65 protein

The physicochemical characteristics of Gp65 were analysed using the GRAVY index (http://www.gravy-calculator.de/index.php) and ProtParam tool (https://web.expasy.org/protparam/) (Gasteiger *et al*., 2005). To study the polarity, percentage accessibility and flexibility profile of the protein, ProtScale server of ExPASy (https://web.expasy.org/protscale/) (Gasteiger *et al*., 2005) was used and the sequence was analysed in a standard nine window scale. The secondary structure of the protein was predicted using PredictProtein Program (https://predictprotein.org/) (Rost *et al*., 2004) and the fold pattern of Gp65 was determined using PFP-FunDSeqE Server (http://www.csbio.sjtu.edu.cn/bioinf/PFP-FunDSeqE/) (Shen and Chou, 2009)

#### 2.1.3 Prediction of nucleotide-binding residues in Gp65 protein

DP-Bind (http://lcg.rit.albany.edu/dp-bind/) (Hwang and Kuznetsov, 2007) and DRNApred tools (http://biomine.cs.vcu.edu/servers/DRNApred/) (Yan and Kurgan, 2017) were used to confirm Gp65 as a DNA-binding protein. To predict the ATP interacting residues in Gp65, the amino acid sequence was analysed using ATPint (http://crdd.osdd.net/raghava/atpint/) (Chauhan *et al*., 2009), which utilizes a support vector machine (SVM) approach. PROVEAN tool (http://provean.jcvi.org/index.php) (Choi and Chan, 2015) was used to analyse the effect of substitution of amino acid residue in the activity of Gp65.

#### 2.1.4 Structural modelling of Gp65 and molecular docking of ATP

Structure of Gp65-AAA ATPase was modelled using a homology/ *ab initio* hybrid tool BHAGEERATH-H (http://www.scfbio-iitd.res.in/bhageerath/bhageerath_h.jsp) (Jayaram *et al*., 2014). 3D scores of the top five predicted structures were validated by analysing Ramachandran plots for each model, using the tool PROCHECK (https://servicesn.mbi.ucla.edu/PROCHECK/) (Laskowski *et al*., 1993) followed by energy minimization using Chiron (https://dokhlab.med.psu.edu/chiron/login.php) (Ramachandran *et al*., 2011). Walker A and Walker B motifs were marked in the selected structure (Model 4) using PyMOL (DeLano, 2002).

To confirm the predicted ATP binding region in Gp65-AAA ATPase, site-specific molecular docking was carried out using AutoDock 1.5.6 tool of MGL software package and AutoDock Vina (Morris *et al*., 2009; Vina, 2010). ATP binding site predicted in this study (using ATPint tool) and reported previously for AAA ATPase enzymes was evaluated as an active site for docking (Ogura and Wilkinson, 2001). Gp65-AAA ATPase 3D structure was prepared using AutoDock tool, which included removal of water molecules and addition of hydrogen bonds. The grid box was set at 66.703, 64.627 and 67.508 at X, Y and Z centres, respectively which included the active site residues Gly21, Pro22, Ser23, Gly24, Ser25, Gly26, Lys27 and Thr28.

The ATP (ligand) SDF file (3D) was downloaded from the PubChem database (https://pubchem.ncbi.nlm.nih.gov/) (Kim *et al*., 2019) and converted to PDB format using BIOVIA Discovery Studio Visualizer tool. To the PDB file of ATP, Gasteiger charges were added, 8 non-polar hydrogens were merged and PDBQT input files were prepared for the receptor and ligand. Molecular docking was performed using Autodock Vina and ten possible conformations were given out as results which were visually rescored using PyMOL.

#### 2.1.5 Functional prediction of Gp65 protein

To predict the putative function of Gp65 protein in *SimranZ1* mycobacteriophage, the protein was analysed using the I-TASSER tool (https://zhanglab.ccmb.med.umich.edu/I-TASSER/) (Zhang, 2008).

### 2.2 Experimental validation of the predicted molecular function of Gp65 protein

#### 2.2.1 Bacterial strains, plasmids and chemicals

*E. coli* DH5 *alpha* and *E. coli* BL21 (DE3) strains were used for cloning and expression studies. Genomic DNA of mycobacteriophage *SimranZ1* was isolated using a standard phenol-chloroform SDS extraction method as described previously (Sinha *et al*., 2020). Restriction enzymes, *Taq* DNA Polymerase and T4 DNA ligase were obtained from New England Biolabs (USA). Plasmid pET28a was kindly gifted by Dr. Ramachandran (IGIB, New Delhi) and for plasmid isolation, the UniPro Plasmid mini kit (India) was used. Phenylmethanesulfonylfluoride (PMSF) and all other analytical grade chemicals were purchased from Himedia chemicals (India). Standard recombinant DNA techniques were used as described elsewhere (Sambrook, 2000).

#### 2.2.2 PCR amplification, cloning and expression of Gp65

Gp65 gene was amplified using *SimranZ1* genomic DNA as a template and gene-specific primers: Gp65 Forward primer 5’-CCG<u>GAATTCA</u>TGAGCCTGTCTTTCAAACC-3’ and Gp65 Reverse primer 5’-CCC<u>AAGCTTT</u>CACTGTTCGGCTCTTTTC-3’ containing EcoRI and HindIII restriction sites, respectively. Phusion®High-Fidelity DNA polymerase was used to amplify the gene.

The amplicon was cloned in the pET28a expression vector and pET28a-Gp65 was transformed in *E.coli* BL21 (DE3) strain. For expression studies, the transformed cells were grown in 10 ml LB medium containing kanamycin (50 µg/ml) for 3-4 hours at 37 °C (O.D._600_ ∼ 0.6). Uninduced cells were aliquoted separately and to the remaining culture, IPTG was added to a final concentration of 0.1 mM. The culture was incubated overnight at 16 °C with shaking and harvested by centrifugation at 8000 r.p.m for 10 minutes. The induced and uninduced cell pellet obtained was suspended in SDS gel loading buffer and electrophoresed on 12% SDS-PAGE. Protein bands were visualized by staining the gel with Coomassie Brilliant Blue R250 (0.05%).

#### 2.2.3 Purification of Gp65 protein

Recombinant Gp65 protein obtained in the soluble fraction was purified by Ni-NTA affinity chromatography. Briefly, the His_6_ tag fusion protein in the soluble form was mixed with 2 ml of Ni-NTA matrix at 4 °C for 2 hours, packed in a column and the flow-through was discarded. The column was washed using the column buffer and the bound protein was eluted using an elution buffer containing imidazole (20 mM Tris pH 7.9, 500 mM NaCl and 200 mM Imidazole). The eluted fractions were analysed using 12% SDS-PAGE and was found to be purified to about 95% homogeneity. The purified protein was dialyzed against storage buffer (50 mM Tris pH 7.9, 150 mM NaCl and 10% Glycerol) and its concentration was estimated by Bradford method (Kruger, 2009).

#### 2.2.4 Western blotting analysis of Gp65

Western blotting of Gp65 was performed by separating the protein sample in a 12% SDS PAGE. The protein was then transferred to a PVDF membrane by using Mini-PROTEAN Tetra Cell (Bio-Rad, United States). The membrane was blocked overnight by using 5% BSA at 4 °C followed by incubation with mouse anti-His6 antiserum (Santacruz Biotech, USA) for 4 hours at a dilution of 1:2000. PBS (1 X) was used to wash the membrane to remove un-bound antibodies. After washing, the membrane was incubated with IgG HRP-conjugated antibody at a dilution of 1:5000 for 2 hours at room temperature. The blot was then developed by using 3, 3’-Diaminobenzidine as the substrate.

#### 2.2.5 Analysis of oligomerization in Gp65

To analyse the oligomeric nature of the protein, purified Gp65 was electrophoresed in 8% non-denaturing PAGE (Native PAGE) at 90 V and the gel was stained with Coomassie blue R250 (0.05%).

To reduce disulphide bridges, the purified protein was treated with reducing Laemmli buffer (50 mM Tris-HCl, pH 7.9, 2% SDS, 10% Glycerol, 0.02% bromophenol blue and 5% beta-mercaptoethanol). Protein in the non-reducing Laemmli buffer (50 mM Tris-HCl, pH 7.9, 2% SDS, 10% Glycerol and 0.02% bromophenol blue) was kept as a control. Samples were incubated for 1 hour at room temperature and separated on a 10% SDS PAGE.

#### 2.2.6 Electron microscopy of Gp65

TEM analysis of Gp65 was performed using JEM-1400Flash at the Advanced Technology Platform Centre, Regional Centre of Biotechnology, India. Briefly, Gp65 was immobilized to glow discharged carbon coated copper grids for 2 minutes and then washed twice with sterile water to remove any unbound proteins. The grid was then stained with 2% Phosphotungstic acid and imaging was performed at 1, 20,000 X.

#### 2.2.7 In-vitro ATPase assay

ATPase activity of Gp65 was measured in a 100 µl reaction volume as described by Rule *et al*., 2016 with minor modifications. The reaction conditions were first optimized by studying the effect of i) different concentrations of the enzyme (0.5-4 µM), co-factor MgCl_2_ (2-10 mM) and substrate ATP (0.2–1 mM), ii) using different buffers (Bis-Tris Propane, Tris-HCl and HEPES maintained at pH 8.5), iii) incubation temperatures (4 °C –55 °C) and iv) incubation period (5–60 minutes). Final enzyme reaction contained 50 mM HEPES/NaCl/Glycerol (HNG) buffer (100 mM HEPES pH 8.5, 65 mM NaCl and 5% glycerol), 3 µM Gp65 protein, 0.4 mM ATP and 6 mM MgCl_2_ with incubation for 30 mins at 37 °C. The reactions were terminated using 12.5 µl of P_i_ ColorlockTM Assay Reagent (Innova Biosciences, U.K) and the absorbance was measured at 630 nm. Net P_i_ released in the hydrolysis of ATP was calculated from the standard phosphate curve. Control reactions contained all the components except the enzyme. The assays were performed in triplicate.

#### 2.2.8 Experimental analysis of Gp65 as a DNA-dependent ATPase activity

To analyse DNA binding property of Gp65, a filter binding assay was performed as described by Peter G. Stockley, 2009 and Yanming Liu *et al*., 2012 with minor modifications. Briefly, to activated PVDF membrane (cut approximately 2 cm in diameter) placed in a 12 well plate, 30 µg of Gp65-AAA ATPase was added. The membrane was air dried and then blocked with 5% BSA and 0.25% Tween 20 for 1 hour at room temperature, followed by three washes in 1 X PBS and 0.05% Tween 20 and incubation in binding buffer (1 X PBS, 150 mM KCl, 6 mM MgCl_2_ and 1 mM DTT) for 30 minutes. Genomic DNA (5 µg) from mycobacteriophage *SimranZ1* was then spotted on the membrane and incubated for 2 hour at 37 °C. Washing with 1 X PBS and 0.05% tween was then performed thrice to remove unbound DNA. The membrane strips were immersed in 100 µL of nuclease free water and then heated at 92 °C for 30 minutes allowing any bound DNA to re-dissolve. Throughout the analysis, PVDF strips blocked with BSA alone served as a control. To validate the success of the experiment, PCR of the eluted mixture using primers for Gp65 gene (as stated in the section 2.2.2) was carried out. The assay was performed in triplicate.

DNA-dependent ATPases are known to hydrolyse ATP maximally in the presence of DNA (Nongkhlaw *et al*., 2009). To validate the DNA-dependent ATPase function of Gp65 assigned through *in silico* analysis in this study, ATPase assay (as described in the section 2.2.4) was carried out in the presence of a fixed concentration (2 µg) of dsDNA The control reaction contained all the components but for the dsDNA. The assay was performed in triplicates.

## 3. Results

### 3.1 *In silico* analysis of Gp65-AAA ATPase from *SimranZ1*

#### 3.1.1 Characterization of Gp65 sequence

Sequence analysis predicted Gp65 to share 99.66% identity with AAA ATPase protein of mycobacteriophages Saal, Demsculpinboyz and Wachhund (https://phagesdb.org/). Interestingly, the sequence also shared 81.75% and 81.66% identity with ATP-binding protein associated with various cellular activities of *Mycobacterium kansasii* and *Mycobacteroides abscessus*, respectively. Analysis of multiple sequence alignment file predicted the presence of Walker A (GXXXXGKT/S, X is any amino acid residue) and Walker B (hhhhhD, h is any hydrophobic residue) motif in Gp65 and homologous sequences which form the essential region for ATP binding in the protein (Figure S1) (Hanson and Whiteheart, 2005).

On phylogenetic analysis, Gp65 was found to share homology with AAA ATPase protein of cluster F1 mycobacteriophage Gumbie, Saal, and Wachhund (Figure S2). Conserved domain analysis predicted AAA_24 domain (Gp65^13-192^, E value: 9.93e-30) and RecA/RadA domain (Gp65^17-113^, E value: 3.83e-06) involved in replication, recombination or repair of DNA (Figure 1) (del Val *et al*., 2019). The predicted domains suggested a possible ATP hydrolysis mediated DNA repair/maintenance role of this protein in *SimranZ1* phage.

**Figure 1:**
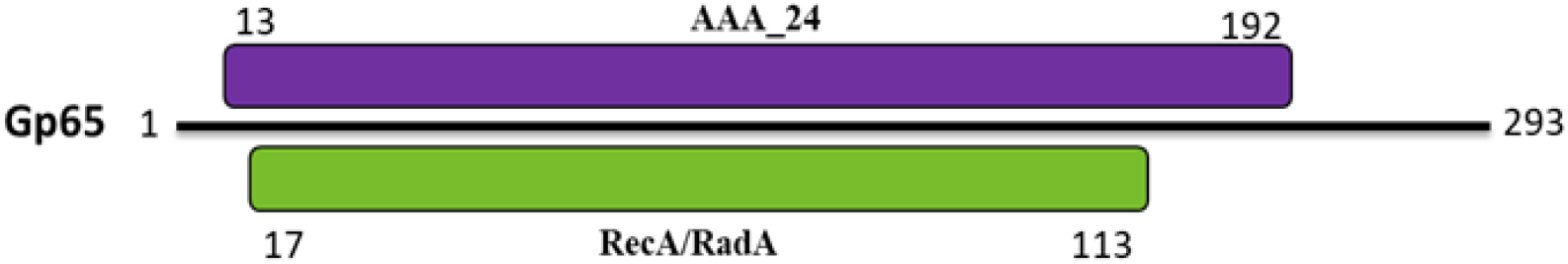
Predicted domain architecture of Gp65-AAA ATPase from SimranZ1. Position of conserved AAA_24 domain involved in ATPase activity is 13aa-192aa and RecA/RadA domain with a putative DNA repair role is 17aa-113aa.

#### 3.1.2 Characterization of AAA ATPase of SimranZ1

*In silico* characterization predicted Gp65-AAA ATPase to have a GRAVY index of −0.363 which classifies the protein to be hydrophilic. The ProtScale server of ExPASy further predicted the high polarity profile of the protein and also suggested the Walker motif regions of the enzyme involved in ATPase activity to be flexible with well accessible residues.

The secondary structure of Gp65 as predicted by the PredictProtein tool, showed the protein to have 42.7% alpha-helix, 21.8% extended strands and 35.5% random coils. Also, evaluation of the protein based on functional domain characterization and sequential evolution information predicted it to have a DNA-binding 3-helical bundle fold, indicating its possible DNA binding role (Contreras-Moreira and Collado-Vides, 2006).

#### 3.1.3 Sequence-based evaluation predicts Gp65’s nucleotide-binding role

DNA binding residues in the sequence were evaluated on a strict consensus-based approach which predicted residues Ala8, Thr9, Arg10, Glu11, Gly24, Ser25, Gly26, Lys27, Thr28, Tyr29, Thr30, Val115, Arg116, Gly117, Asn118, Thr119, Phe120, Arg127, Pro128, Asp129 and Glu130 as DNA binding regions in the protein. To analyse the ATP binding role of Gp65, the protein sequence was evaluated using a sequence-based nucleotide-binding residue prediction tool which predicted the Walker A motif of the protein to constitute ATP binding residues against a threshold of 0.6. Additionally, a PROVEAN tool score of −6.583 and −3.433 (against a standard threshold of −2.5) validated the key role of Lys27 and Asp104 residues in Walker A and Walker B motif, respectively in the function of Gp65 protein.

#### 3.1.4 Structural modelling and docking of Gp65-AAA ATPase

3D structure of Gp65-AAA ATPase was built based on a homology/ *ab initio* prediction software and the results were screened by Ramachandran plot (Figure 2. A). The predicted model had 88.6% residues in the most favoured region and 11% residues in the additional allowed region (Figure 2. B). The structure was further refined and the conserved Walker A and Walker B motif were marked.

**Figure 2:**
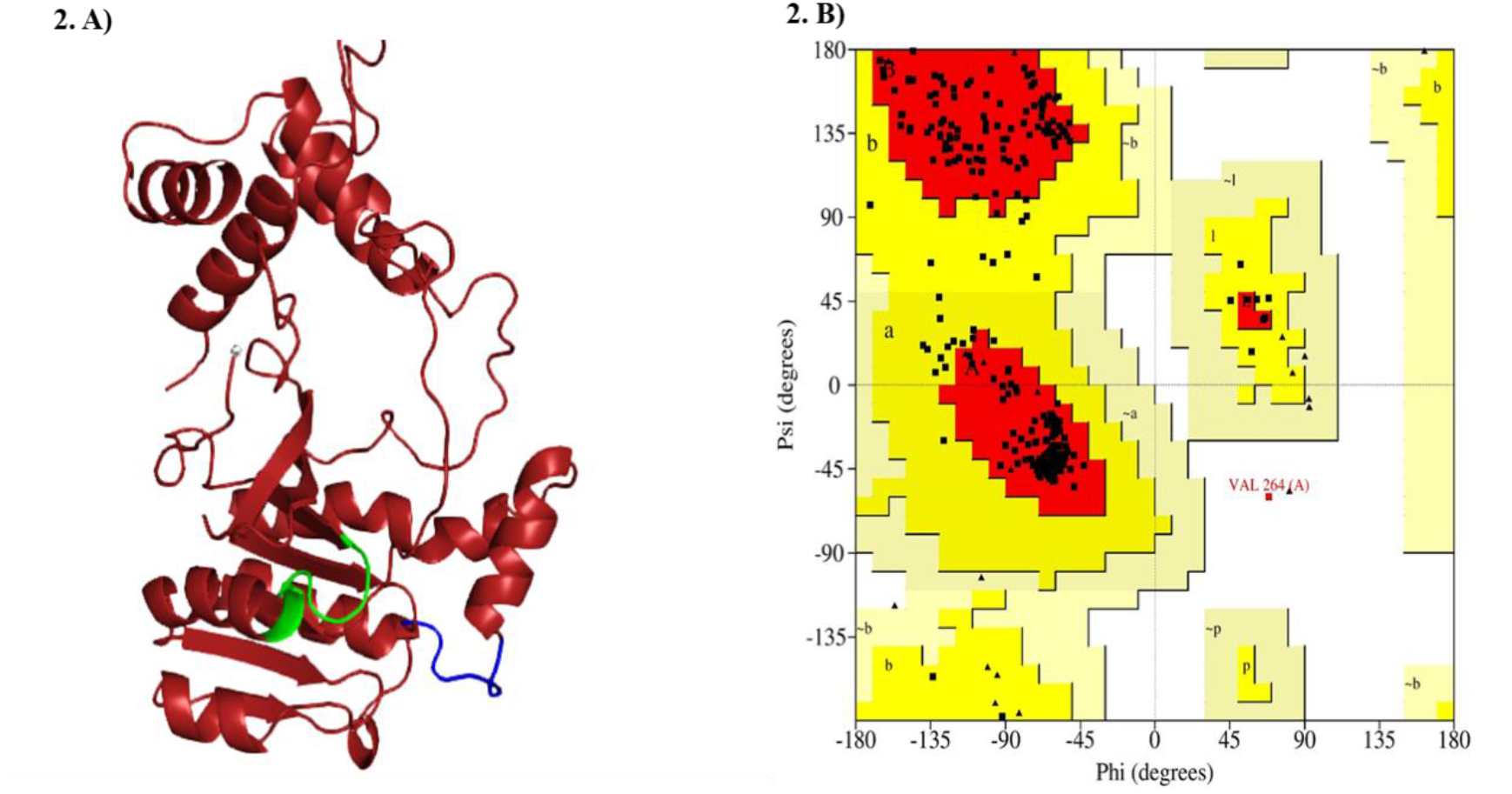
Predicted three-dimensional structure of Gp65-AAA ATPase from *SimranZ1* mycobacteriophage built using homology/*ab initio* hybrid tool (BHAGEERATH-H) and validation by Ramachandran plot. **A)** Highlighted in green is the conserved Walker A motif and highlighted in blue is the conserved Walker B motif which binds to Mg^2^. **B)** Ramachandran plot of 3D structure of Gp65-AAA ATPase indicates 88.6% and 11% residues in the most favoured and additional allowed region respectively and 0.4% residues in the outlier region.

Molecular docking of Gp65-AAA ATPase was performed to examine the interaction of Walker A motif with ATP. On visual rescoring the best conformation using the lowest binding energy (Figure 3. A), we found β and γ phosphate groups of ATP to interact with Gly21 and Ser23 residues in the Walker A motif with a binding affinity of −7.0 kcal/mol (Figure 3. B). This indicates Walker A of Gp65 to be involved in nucleotide-binding and ATPase activity.

**Figure 3:**
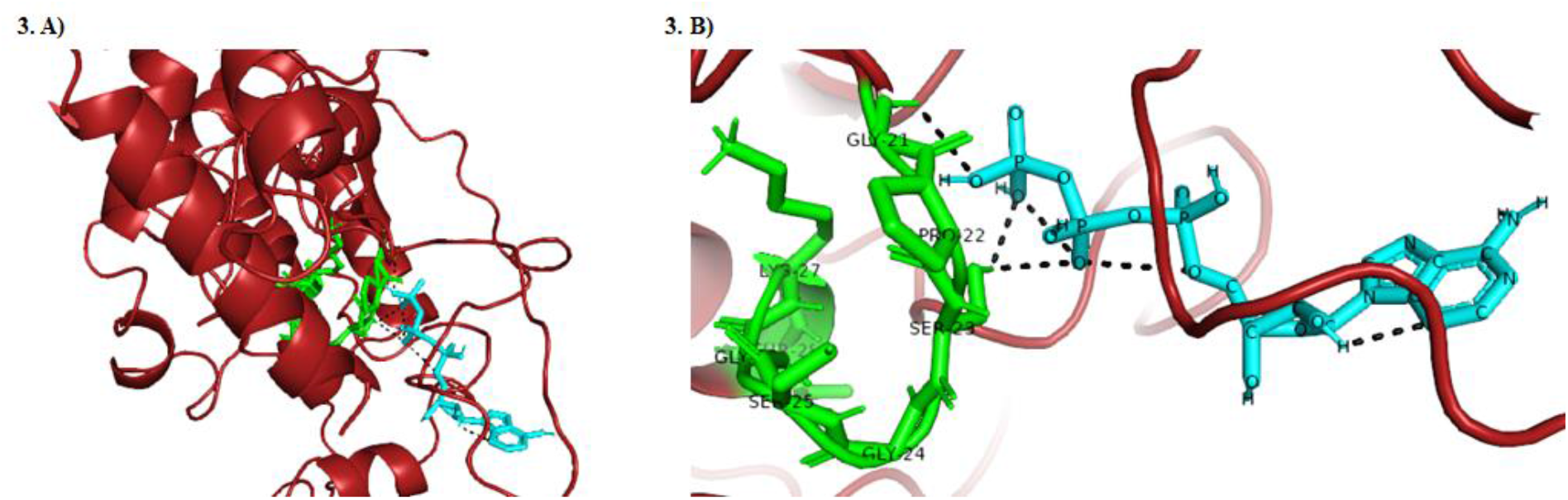
Molecular docking of Gp65-AAA ATPase with ATP. **(A)** Helix-loop-helix structure of Gp65-AAA ATPase shows ATP binding. **(B)** Bird’s eye view of Walker A motif (in green) shows Gly21 and Ser23 to interact (in black) with phosphate groups of ATP (in blue) with a binding affinity of −7 kcal/mol.

#### 3.1.5 In silico prediction of Gp65 function in the SimranZ1 phage

To ascertain the role of Gp65 in *SimranZ1*, the protein was evaluated using the I-TASSER software. The tool predicted Gp65 to be involved in a DNA repair mechanism having a DNA-dependent ATPase role.

### 3.2 Experimental validation of putative function of Gp65

#### 3.2.1 Cloning, expression and purification of Gp65-AAA ATPase

Gp65 gene from *SimranZ1* was amplified by PCR using gene-specific primers (Figure S3) and cloned in pET28a vector. The pET28a-Gp65 plasmid was then transformed into *E.coli* BL21 (DE3) and the recombinant protein (33 kDa) was overexpressed and purified using Nickel-NTA affinity chromatography to 95% homogeneity (Figure 4: A). The purified protein was confirmed by a Western blot using anti-His IgG antibodies (Figure 4: B).

**Figure 4:**
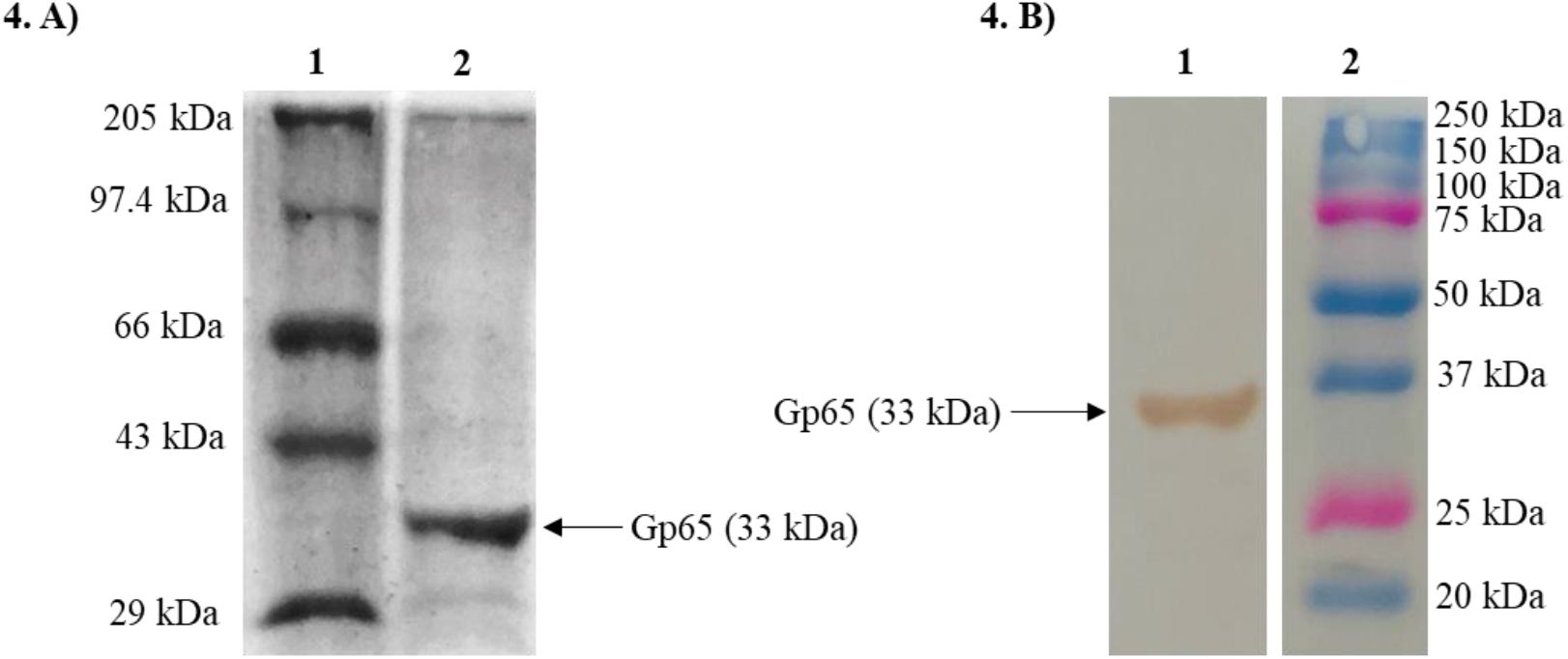
SDS PAGE and Western Blotting of Gp65. **(A)** SDS-PAGE (12%) analysis of purified Gp65-AAA ATPase. Lane 1: Protein molecular mass standards. Lane 2: Purified recombinant protein (33 kDa). **(B)** Western blotting analysis of N-terminal His-tagged Gp65. Lane 1: N-terminal His-tagged Gp65 protein. Lane 2: Protein molecular mass standards.

#### 3.2.2 Gp65-AAA ATPase is a hexameric protein

Native PAGE of purified Gp65 demonstrated that the protein assembled into various oligomeric states (Figure 5: A). The hexameric nature of Gp65 was also confirmed through TEM imaging at 1, 20,000 X. The protein in its hexameric state has a diameter of approximately 8.84 nm consisting of a central pore of diameter 1.9 nm (Figure 5: B).

**Figure 5:**
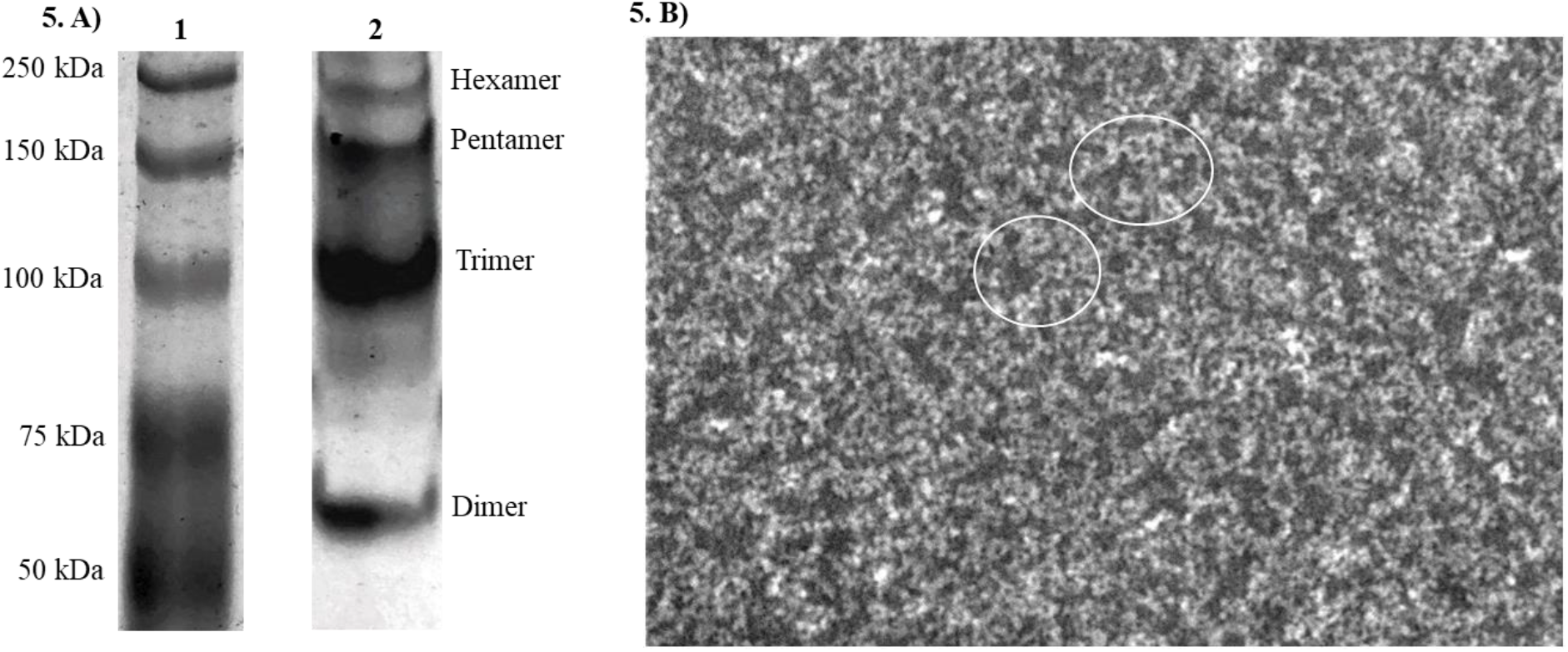
Analysis of oligomerization of Gp65. **A)** 8% Native PAGE analysis of oligomeric forms of the purified Gp65 protein. Lane 1: Protein molecular mass standards. Lane 2: Hexameric, Pentameric, Trimeric and Dimeric forms of Gp65-AAA ATPase. **B)** TEM analysis of Gp65-AAA ATPase. Within the white circles are seen the hexameric forms of the protein.

#### 3.2.3 Gp65-AAA ATPase’s oligomeric forms are held by disulphide bonds

To analyse whether the oligomeric states of Gp65 were formed by covalent bonding (disulphide bridges) between the individual monomers, we carried out 10% SDS PAGE of the purified protein in the presence and absence of a reducing agent (beta-mercaptoethanol). Under reducing conditions, protomer of Gp65 (33 kDa) was obtained (Figure 6: A) while under non-reducing conditions, Gp65 migrated in its monomeric, dimeric, trimeric and tetrameric forms in the PAGE (Figure 6: B).

**Figure 6:**
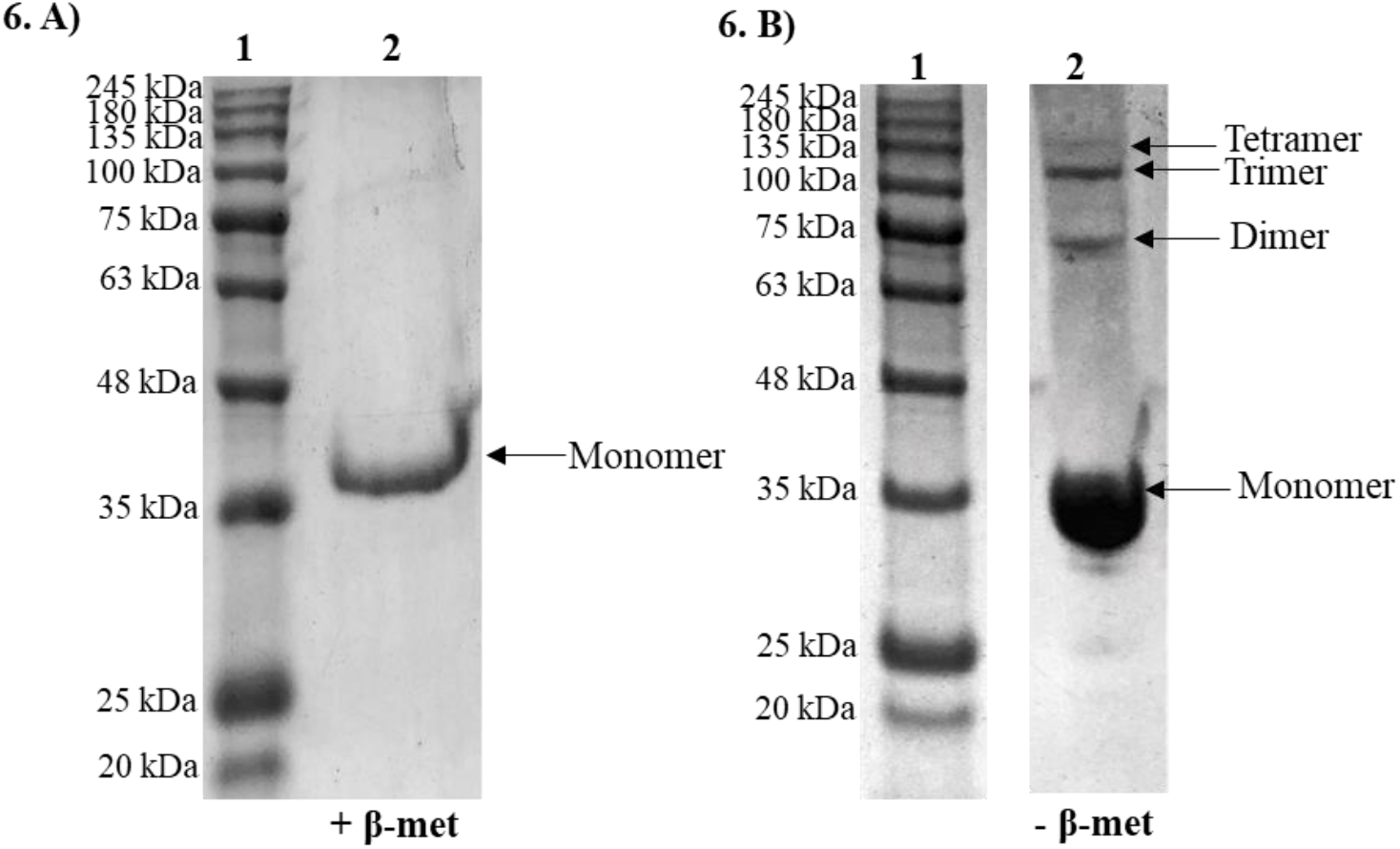
SDS-PAGE analysis of disulphide bonds in Gp65 oligomers. **A)** Gp65 in the presence of reducing agent (β-mercaptoethanol). Lane 1: Protein molecular mass standards. Lane 2: Migration of Gp65 as a monomer in the presence of β-mercaptoethanol. **B)** Gp65 in the absence of reducing agents. Lane 1: Protein molecular mass standards. Lane 2: Migration of Gp65 in its Tetrameric, Trimeric, Dimeric and Monomeric forms.

#### 3.2.4 Optimization of ATPase assay

ATPase activity of Gp65 was optimized under different conditions (Figure 7, 8 and 9) and was found to have an activity of 0.537 (± 0.017) nmol Pi/min at 3 μM enzyme concentration when incubated for 30 minutes at 37 °C in HEPES pH 8.5 buffer. The V_max_, K_m_ and k_cat_ of the enzyme was found to be 0.42 mM/ min, 0.098 mM and 0.0023 s^-1^, respectively (Table 1).

**Figure 7:**
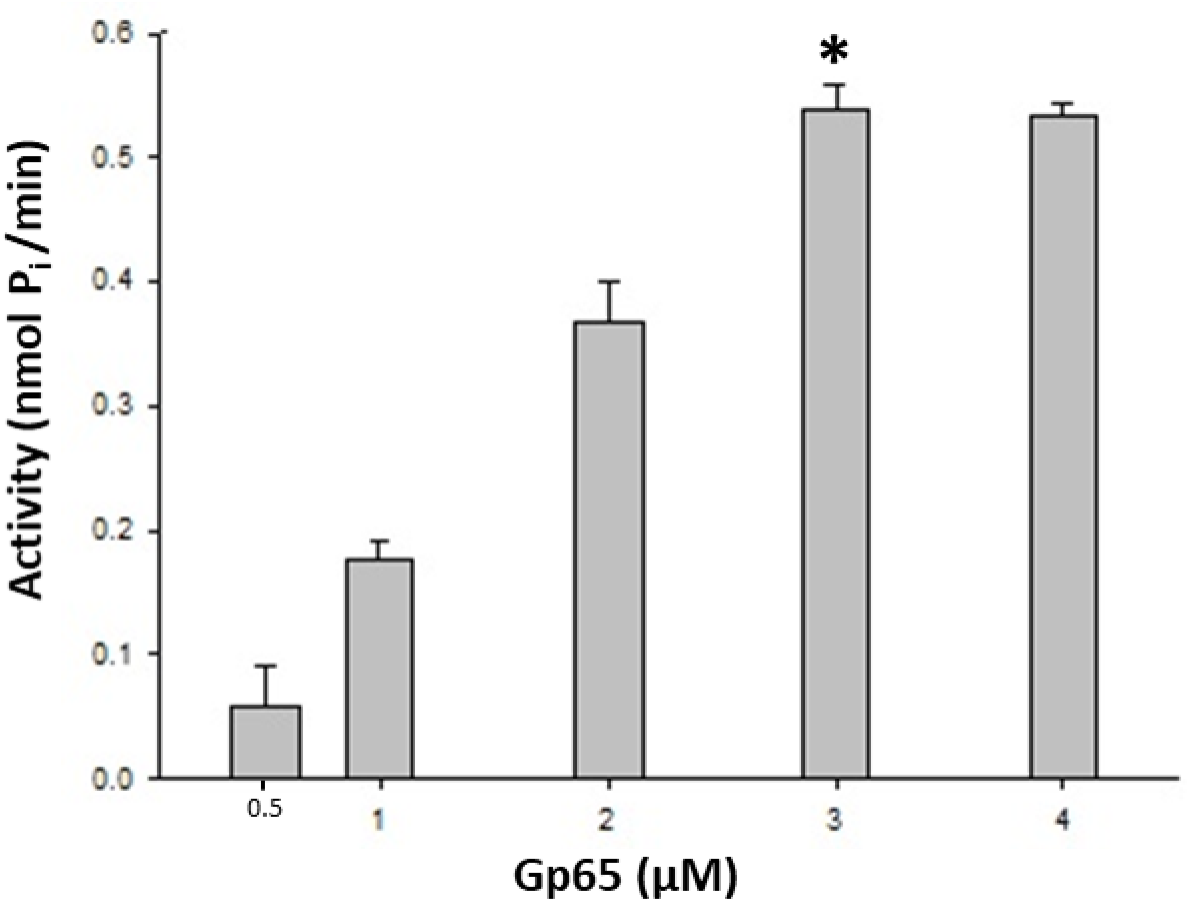
Activity of Gp65-AAA ATPase at various concentrations. X-axis represents the concentration of Gp65 enzyme (µM); Y-axis represents the net activity of enzyme (nmol P_i_/min). Data depicted are the mean ±S.E values observed from three independent experiments. ‘*’ represents the concentration of Gp65 used in the subsequent assays.

**Figure 8:**
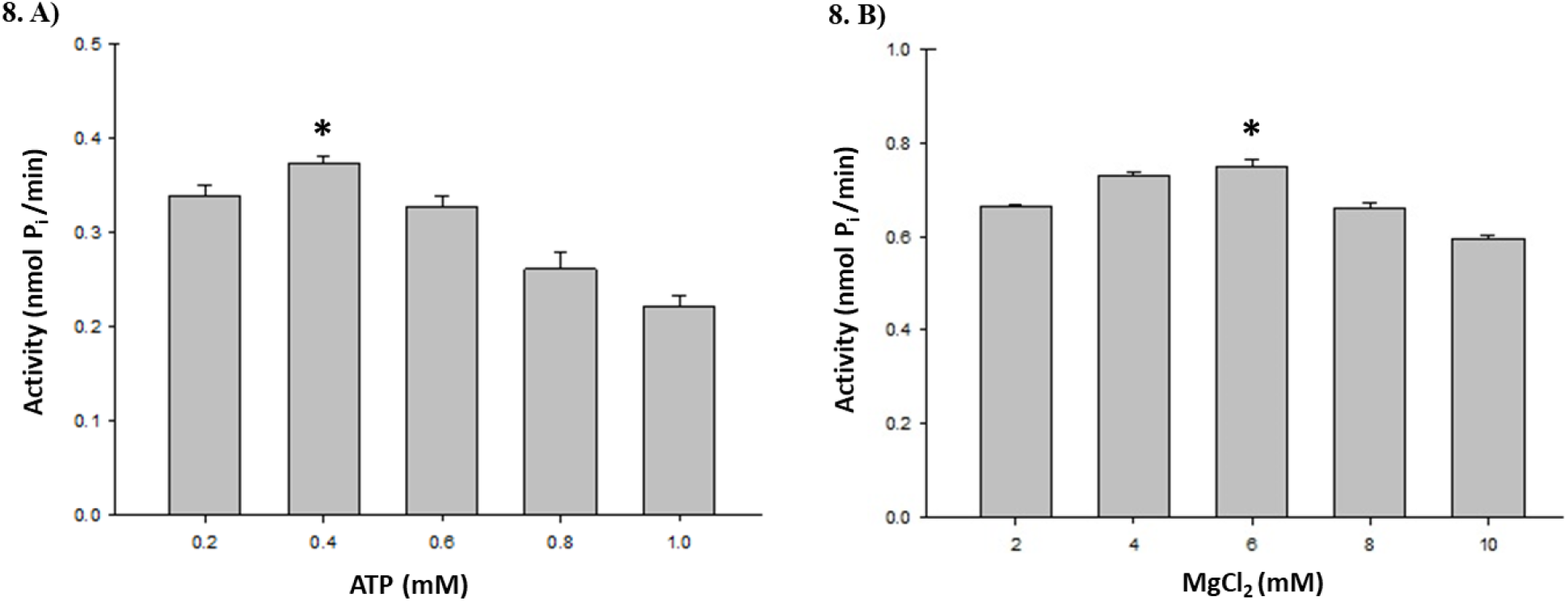
Activity of Gp65-AAA ATPase at various concentrations of ATP and MgCl_2_. **(A)** X-axis represents the concentration of ATP (mM); Y-axis represents the activity (nmol P_i_/min) of Gp65-AAA ATPase. **(B)** X-axis represents the concentration of MgCl_2_ (mM); Y-axis represents the activity (nmol P_i_/min) of Gp65-AAA ATPase. Data depicted are the mean ±S.E. values observed from three independent experiments. ‘*’ represents the final concentration of ATP and MgCl_2_ used in the assays.

**Figure 9:**
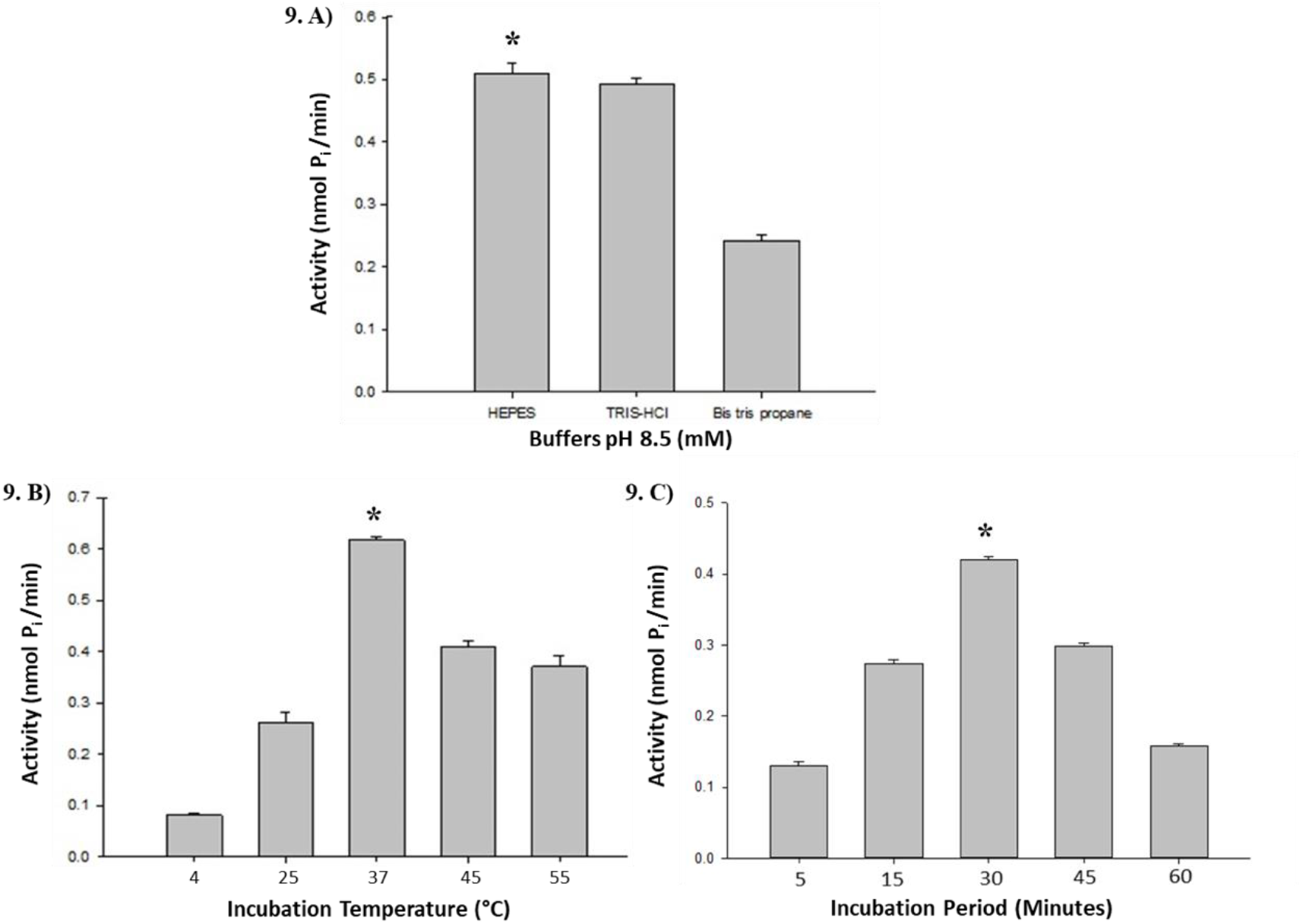
Activity of Gp65-AAA ATPase at different buffers, temperature and time period. **(A)** X-axis represents different buffers at pH 8.5; Y-axis represents the activity (nmol P_i_/min) of Gp65-AAA ATPase. **(B)** X-axis represents the incubation temperature (°C); Y-axis represents the activity (nmol P_i_/min) of Gp65-AAA ATPase. Optimum enzyme activity was observed at 37°C. **(C)** X-axis represents the incubation period (minutes); Y-axis represents the activity (nmol P_i_/min) of Gp65-AAA ATPase. Data depicted are the mean ±S.E values observed from three independent experiments. ‘*’ represents the incubation temperature, time period and buffer conditions used in the assay.

**Table 1:**
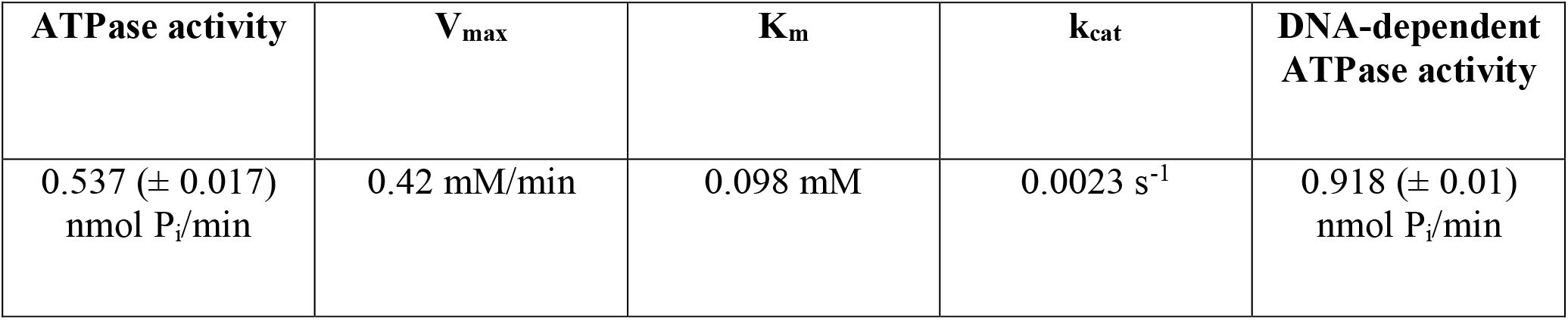
Kinetic parameters for Gp65-AAA ATPase activity

| **ATPase activity** | **$V_{\max}$** | **$K_m$** | **$k_{\text{cat}}$** | **DNA-dependent ATPase activity** |
| --- | --- | --- | --- | --- |
| 0.537 (± 0.017)<br>nmol $P_i$/min | 0.42 mM/min | 0.098 mM | 0.0023 s<sup>-1</sup> | 0.918 (± 0.01)<br>nmol $P_i$/min |

1. A)

#### 3.2.5 Gp65 is a DNA binding protein

A modified version of the filter binding assay was performed to demonstrate that Gp65 binds with double stranded DNA. The DNA eluted after washing the PVDF membrane strips were subjected to PCR with primers for Gp65 gene. Amplification was observed in the eluted fraction of the experimental sample (Gp65 protein) and not in the case of BSA which served as a negative control (Figure S4).

#### 3.2.6 DNA stimulates Gp65’s ATPase activity

The DNA-dependent ATPase assay of Gp65 was carried out in the presence of dsDNA (2 μg). The activity of the enzyme was found to be 0.918 (± 0.01) nmol P_i_/min (Table 1), showing a relative increase of 70.9% in the activity (Figure 10).

**Figure 10:**
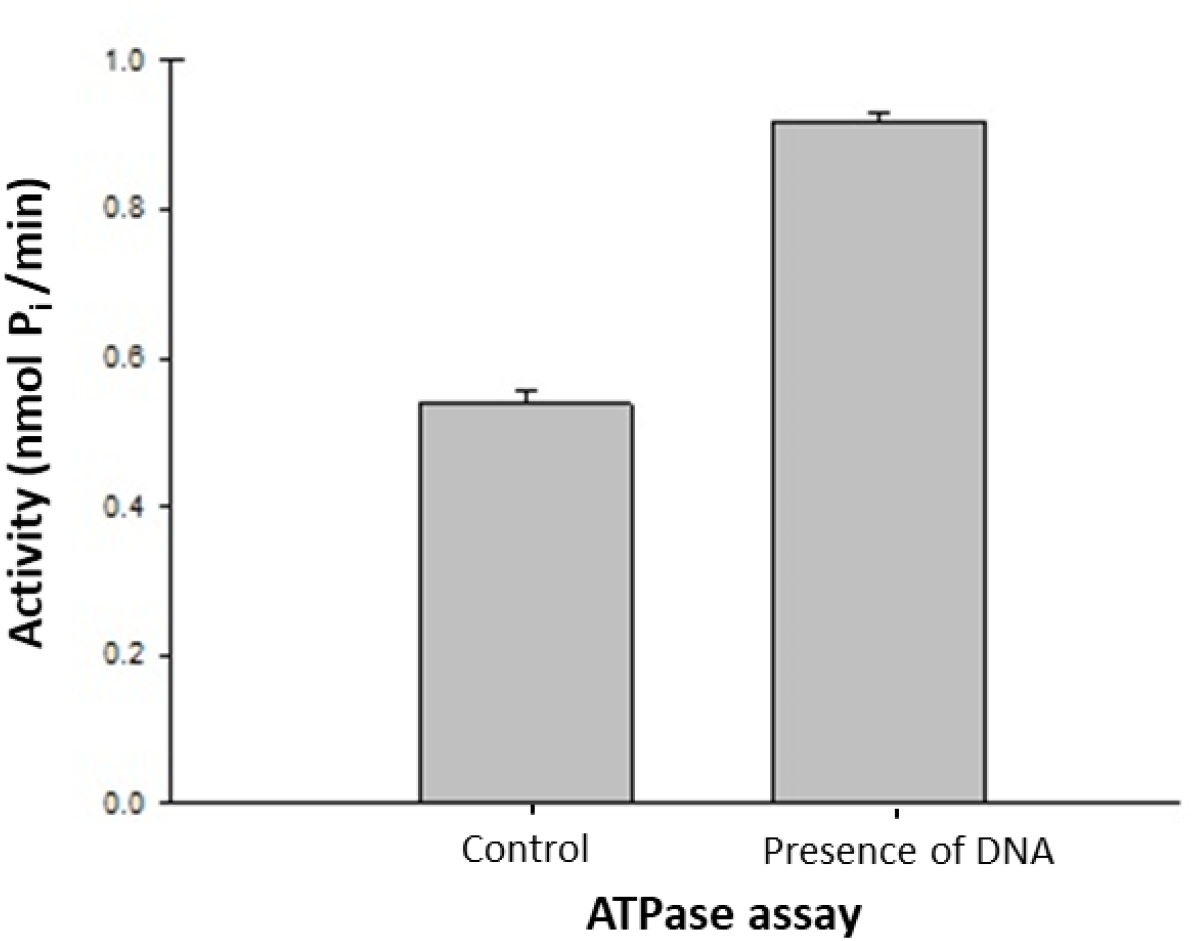
DNA-dependent ATPase assay of Gp65. X-axis represents the ATPase assay in the absence of dsDNA (control) and in the presence of dsDNA (2 μg). Y-axis represents the net activity (nmol P_i_/min) in each assay. Control reaction contained all the components except for dsDNA. Data depicted are the mean ±S.E values observed from three independent experiments.

## 4. Discussion

The infections caused by *Mycobacterium* spp. and an increase in the rise of drug-resistant strains poses an enormous threat to humankind. In the absence of new drugs, well characterized lytic mycobacteriophages are being considered as promising candidates to offer an effective anti-mycobacterial solution (Hatfull, 2014). Isolation of novel mycobacteriophages, their genomic characterization and understanding the function of their encoded proteins therefore is of paramount importance. However, the fact that the function of a large part of phage genomes is unknown can be a potential impediment, which can thwart the acceptance of bacteriophages as therapeutics. Hence, functional annotation followed by experimental evidence can lead to greater confidence in ‘knowing’ the phages and can also contribute to the existing knowledge on diverse roles phage-encoded proteins can play (Skurnik and Strauch, 2006).

According to the Actinobacteriophage database, 182 phages out of the sequenced mycobacteriophage genomes belong to cluster F. Based on the similarity of the nucleotide sequences, the F cluster is sub-divided into 5 sub-clusters and the average genome size is 57, 397 bp. While most of the encoded proteins of the F cluster phages are reported to be present in mycobacteriophages from other clusters, we found Pham 9410 to be present exclusively in F cluster phages and in 7 Arthrobacter and 3 Gordonia phages (https://phagesdb.org/). Its biological function in the phage is yet to be demonstrated though.

AAA ATPase enzymes (belonging to the P-loop NTPase superfamily) from bacteriophages have reported to be the molecular motors, which catalyse the packaging of the viral genome in an energy-dependent manner (Lin *et al*., 2017). In *SimranZ1*, Gp2 and Gp3 are predicted to encode for large and small subunits of terminase, respectively (unpublished data). We studied the structure and function of Gp65 which codes for another AAA ATPase in this phage. Molecular docking of the predicted structure with ATP showed Gly21 and Ser23 residues from Walker A motif to interact with phosphate groups of ATP with a binding energy of −7 kcal/mol, hence suggesting an ATPase functionality. Further bioinformatics analysis of the protein sequence and the domain suggested Gp65 to be a DNA-dependent ATPase, involved in DNA repair and maintenance.

Typically, AAA ATPase proteins having enzymatic activity are known to arrange into hexameric ring complexes (Snider *et al*., 2008), which was observed in the case of Gp65 too as evident by native PAGE and TEM analysis. The central pores of AAA ATPase are responsible for the enzymatic activity of the protein and in the case of Gp65-AAA ATPase (pore size of 1.9 nm) which exhibited DNA-binding activity, we speculate it to be involved in an energy mediated repair of DNA. Protein oligomerization can occur naturally through non-covalent bonds/interactions or through covalent bonds.

Native PAGE analysis of Gp65 in reducing and non-reducing environments revealed the presence of disulphide linkages between the monomeric forms in the oligomerized protein.

The predicted ATPase activity of Gp65 was validated by *in vitro* assay. Its DNA binding property was analysed using a modified version of the filter binding assay and DNA dependent ATPase activity was measured by the addition of dsDNA in the ATPase assay. This the biochemical characterization and the prediction of putative RecA/RadA domain in the protein throws light on the possible role(s) AAA ATPase can have in the life cycle of mycobacteriophages.

## Conclusion

Gp65 from an F1 sub-cluster mycobacteriophage ‘*SimranZ1*’ is a DNA-dependent ATPase with putative DNA repair function. This study consists of bioinformatics analysis of Gp65 which includes *in silico* characterization of the protein, structure prediction and docking studies with ATP, followed by cloning, expression and purification of the recombinant protein and experimental validation of the predicted functions using an *in vitro* microtiter assay. It will be pertinent to further explore the essentiality of ATP-dependent DNA repair/maintenance role of Gp65 in the F cluster mycobacteriophages.

## Acknowledgements

We thank Science and Engineering Research Board (SERB), Department of Science and Technology (DST) (Grant number: EMR/2017/004051), India for supporting the project. We thank Ms Jyoti Rani, Ph.D. and Ms Ritu Arora, research scholars at Acharya Narendra Dev College for the extended assistance in carrying out some of the experiments (molecular docking and cloning of Gp65) and Dr. Eniyan Kandasamy, Senior Postdoctoral Fellow at International Centre for Genetic Engineering and Biotechnology, New Delhi for proofreading the manuscript. We thank the Principal, Acharya Narendra Dev College for Education in a Lively Innovative Training Environment (ELITE) Scholarship and for the infrastructural support.

## Availability of data and material

Data available within the article or its supplementary materials.

## Supplementary Information

**Figure S1:**
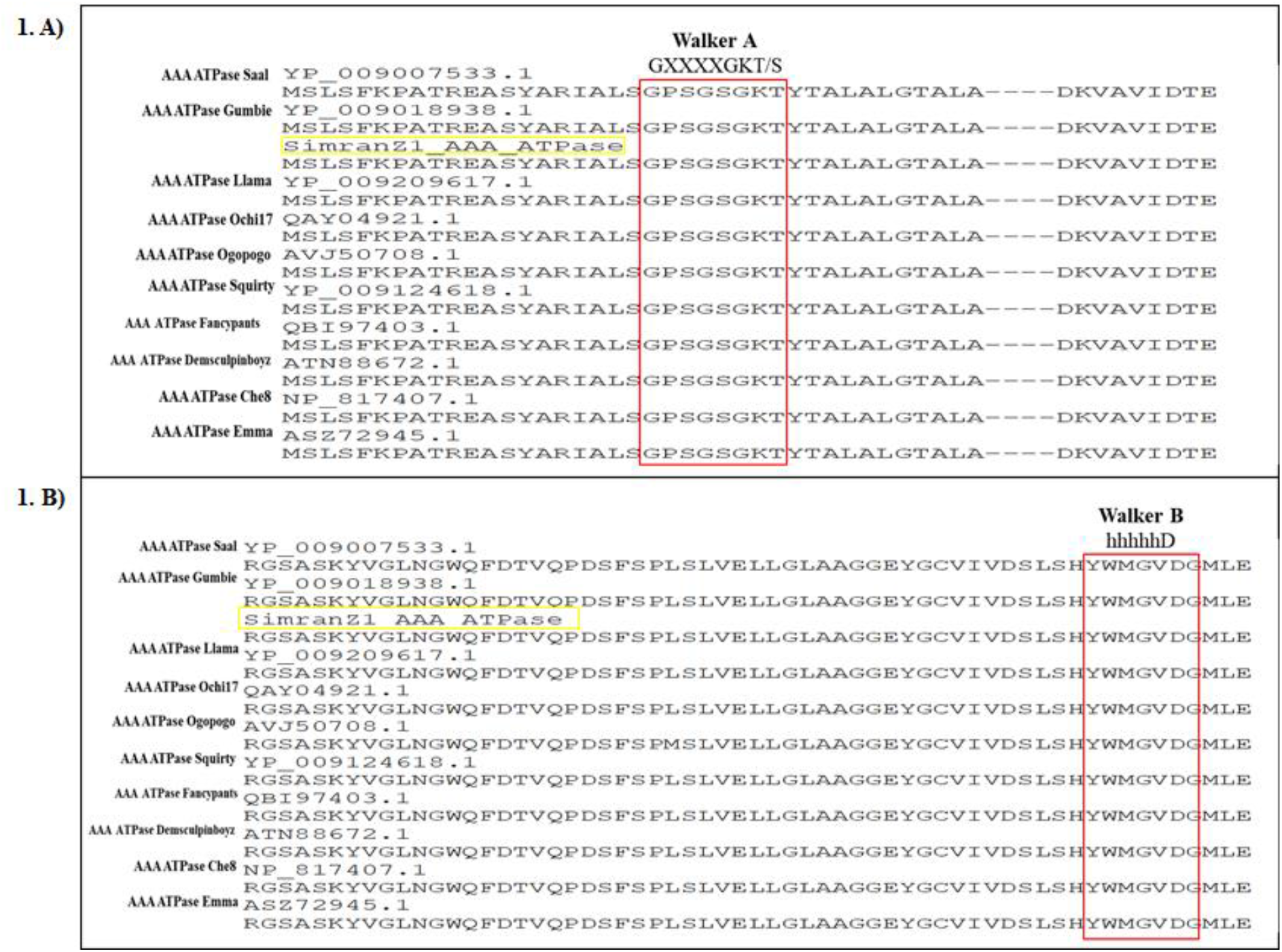
Multiple sequence alignment of AAA ATPase from mycobacteriophages denotes the conserved walker motifs (highlighted in red). (**A)** Conserved Walker A motif (GXXXXGKT/S, where ‘X’ is an amino acid residue) is the phosphate and nucleotide-binding region of the AAA ATPase. (**B)** Shows the conserved Walker B motif (hhhhhD, where ‘h’ is any hydrophobic amino acid residue) of AAA ATPase that binds to Mg^2+^ and the acidic residues are involved in ATPase activity.

**Figure S2:**
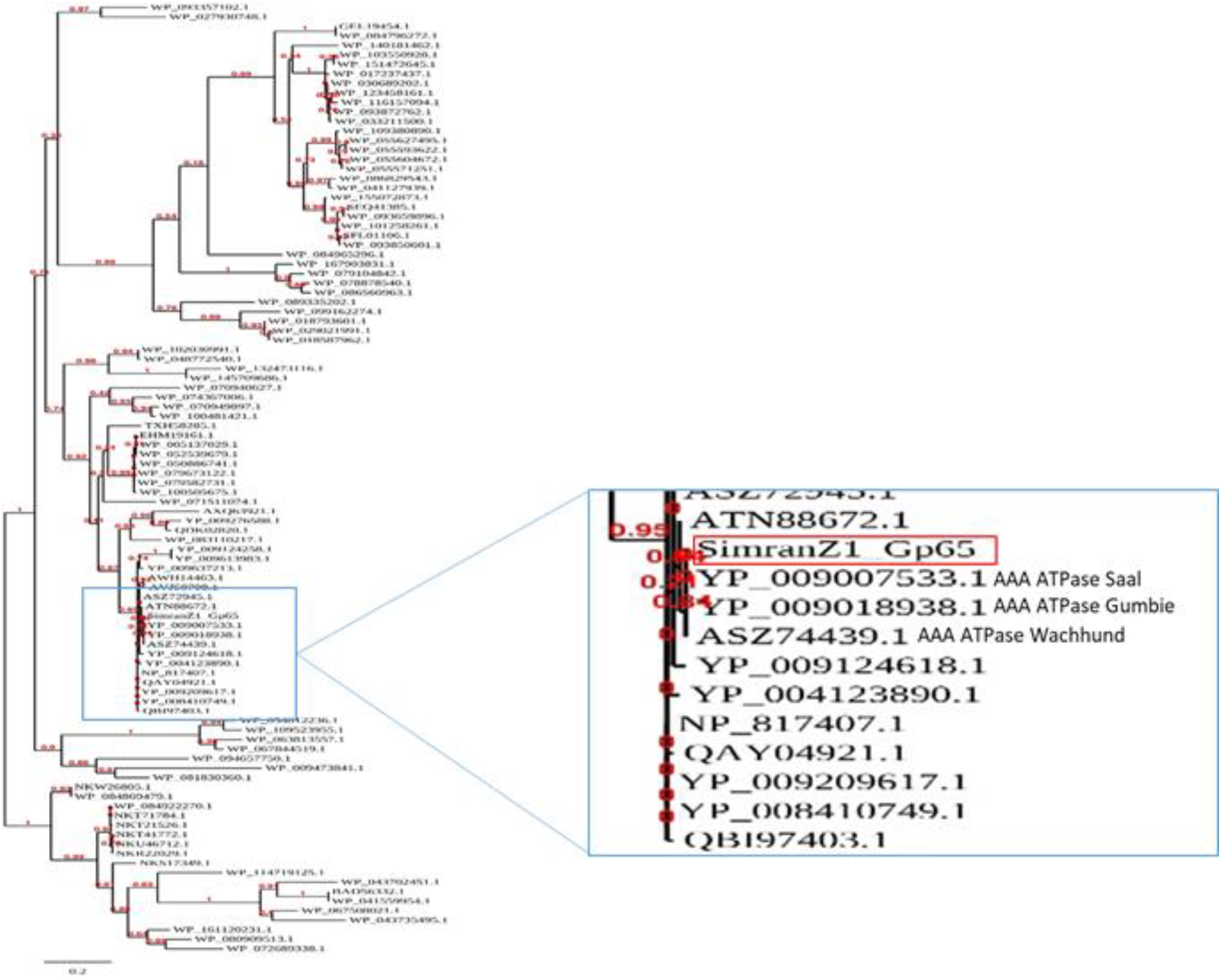
Phylogenetic analysis of Gp65. Homology study through phylogenetic tree analysis of *SimranZ1* Gp65 sequence (highlighted in red) shows homology with AAA ATPase sequence of mycobacteriophage Saal, Gumbie Wachhund and with ATP binding proteins of *Mycobacterium kansasii* and *Mycobacteroides abscessus*.

**Figure S3:**
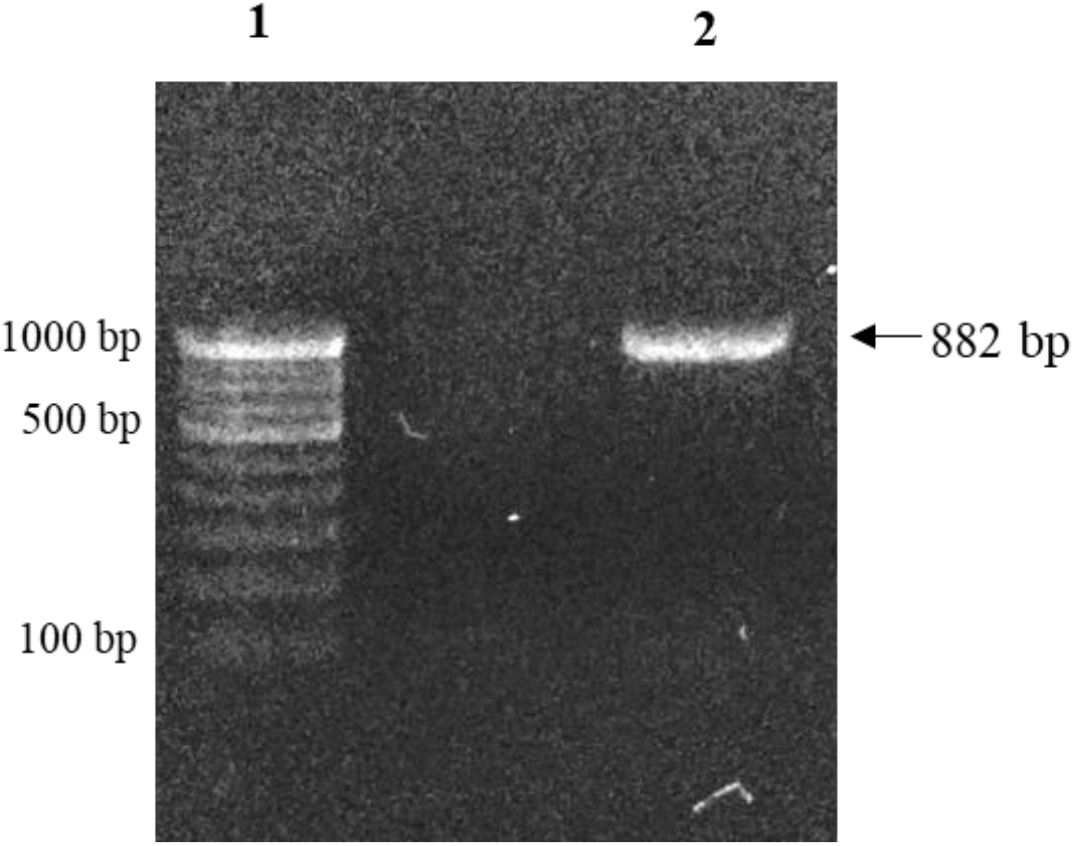
PCR amplification of Gp65 gene. Gp65 gene (882 bp) was amplified by PCR using *SimranZ1* genomic DNA as a template. Lane 1: 100 bp DNA Ladder, Lane 2: Gp65 PCR product.

**Figure S4:**
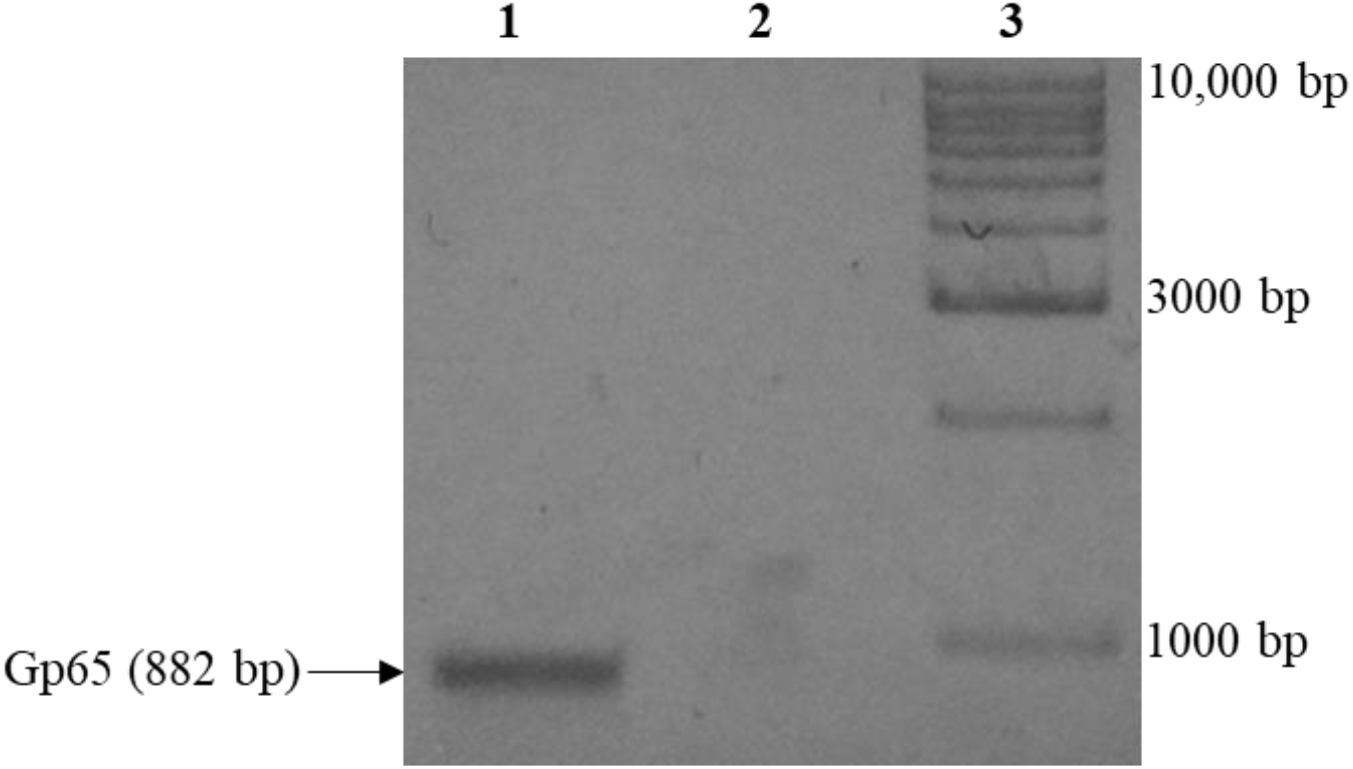
Filter binding assay of Gp65. Lane 1: Amplification of Gp65 gene indicates the presence of DNA in the eluted fraction of experimental sample containing Gp65 protein. Lane 2: No amplification was observed in the negative control containing BSA. Lane 3: 1 kb DNA ladder.

## References

Altschul, S. F., Gish, W., Miller, W., Myers, E. W., & Lipman, D. J. (1990). Basic local alignment search tool. Journal of molecular biology, 215(3), 403 –410.

Bajpai, U., Mehta, A. K., Eniyan, K., Sinha, A., Ray, A., Virdi, S.…, & Chauhan, G. (2018). Isolation and characterization of bacteriophages from India, with lytic activity against Mycobacterium tuberculosis. Canadian journal of microbiology, 64(7), 483 –491.

Chauhan, J. S., Mishra, N. K., & Raghava, G. P. (2009). Identification of ATP binding residues of a protein from its primary sequence. BMC bioinformatics, 10(1), 434.

Choi, Y., & Chan, A. P. (2015). PROVEAN web server: a tool to predict the functional effect of amino acid substitutions and indels. Bioinformatics, 31(16), 2745–2747.

Contreras-Moreira, B., & Collado-Vides, J. (2006). Comparative footprinting of DNA-binding proteins. Bioinformatics, 22(14), e74–e80.

Dedrick, R. M., Guerrero-Bustamante, C. A., Garlena, R. A., Russell, D. A., Ford, K., Harris, K.…, & Hatfull, G. F. (2019). Engineered bacteriophages for treatment of a patient with a disseminated drug-resistant Mycobacterium abscessus. Nature medicine, 25(5), 730–733.

del Val, E., Nasser, W., Abaibou, H., & Reverchon, S. (2019). RecA and DNA recombination: a review of molecular mechanisms. Biochemical Society Transactions, 47(5), 1511–1531.

DeLano, W. L. (2002). PyMOL.

Dereeper, A., Guignon, V., Blanc, G., Audic, S., Buffet, S., Chevenet, F.…, & Claverie, J. M. (2008). Phylogeny. fr: robust phylogenetic analysis for the non-specialist. Nucleic acids research, 36(Suppl_2), W465–W469.

Gasteiger, E., Hoogland, C., Gattiker, A., Wilkins, M. R., Appel, R. D., & Bairoch, A. (2005). Protein identification and analysis tools on the ExPASy server. In The proteomics protocols handbook (pp. 571–607). Humana press.

Hanson, P. I., & Whiteheart, S. W. (2005). AAA+ proteins: have engine, w ill work. Nature reviews Molecular cell biology, 6(7), 519 –529.

Hatfull, G. F. (2014). Mycobacteriophages: windows into tuberculosis. PLoS pathogens, 10(3).

Hatfull, G. F. (2019). Mycobacteriophages. Gram-Positive Pathogens, 1029–1055.

Hatfull, G. F., Pedulla, M. L., Jacobs-Sera, D., Cichon, P. M., Foley, A., Ford, M. E.…, & Namburi, S. (2006). Exploring the mycobacteriophage metaproteome: phage genomics as an educational platform. PLoS genetics, 2(6).

Hilbert, B. J., Hayes, J. A., Stone, N. P., Duffy, C. M., Sankaran, B., & Kelch, B. A. (2015). Structure and mechanism of the ATPase that powers viral genome packaging. Proceedings of the National Academy of Sciences, 112(29), E3792 –E3799.

Hwang, S., Gou, Z., & Kuznetsov, I. B. (2007). DP-Bind: a web server for sequence-based prediction of DNA-binding residues in DNA-binding proteins. Bioinformatics, 23(5), 634 –636.

Jayaram, B., Dhingra, P., Mishra, A., Kaushik, R., Mukherjee, G., Singh, A., & Shekhar, S. (2014). Bhageerath-H: a homology/ab initio hybrid server for predicting tertiary structures of monomeric soluble proteins. BMC bioinformatics, 15(S16), S7.

Kim, S., Chen, J., Cheng, T., Gindulyte, A., He, J., He, S.…, & Zaslavsky, L. (2019). PubChem 2019 update: improved access to chemical data. Nucleic acids research, 47(D1), D1102–D1109.

Kruger, N. J. (2009). The Bradford method for protein quantitation. In The protein protocols handbook (pp. 17–24). Humana Press, Totowa, NJ.

Laskowski, R. A., MacArthur, M. W., Moss, D. S., & Thornton, J. M. (1993). PROCHECK: a program to check the stereochemical quality of protein structures. Journal of applied crystallography, 26(2), 283–291.

Lin, S., Alam, T. I., Kottadiel, V. I., VanGessel, C. J., Tang, W. C., Chemla, Y. R., & Rao, V.B. (2017). Altering the speed of a DNA packaging motor from bacteriophage T4. Nucleic acids research, 45(19), 11437–11448.

Liu, Y., Wang, C., Li, F., Shen, S., Tyrrell, D. L. J., Le, X. C., & Li, X. F. (2012). DNase- mediated single-cycle selection of aptamers for proteins blotted on a membrane. Analytical chemistry, 84(18), 7603–7606.

Madeira, F., Park, Y. M., Lee, J., Buso, N., Gur, T., Madhusoodanan, N., & Lopez, R. (2019). The EMBL-EBI search and sequence analysis tools APIs in 2019. Nucleic acids research, 47(W1), W636–W641.

Marchler-Bauer, A., Lu, S., Anderson, J. B., Chitsaz, F., Derbyshire, M. K.,DeWeese-Scott, C. & Gwadz, M. (2010). CDD: a Conserved Domain Database for the functional annotation of proteins. Nucleic acids research, 39(Suppl_1), D225 –D229.

McNerney, R. (1999). TB: the return of the phage. A review of fifty years of mycobacteriophage research. The International Journal of Tuberculosis and Lung Disease, 3(3), 179–184.

Morris, G. M., Huey, R., Lindstrom, W., Sanner, M. F., Belew, R. K., Goodsell, D. S., & Olson, A. J. (2009). AutoDock4 and AutoDockTools4: Automated docking with selective receptor flexibility. Journal of computational chemistry, 30(16), 2785–2791.

Nongkhlaw, M., Dutta, P., Hockensmith, J. W., Komath, S. S., & Muthuswami, R. (2009). Elucidating the mechanism of DNA-dependent ATP hydrolysis mediated by DNA-dependent ATPase A, a member of the SWI2/SNF2 protein family. Nucleic acids research, 37(10), 3332–3341.

Ogura, T., & Wilkinson, A. J. (2001). AAA+ superfamily ATPases: common structure –diverse function. Genes to Cells, 6(7), 575–597.

Pope, W. H., Bowman, C. A., Russell, D. A., Jacobs-Sera, D., Asai, D. J., Cresawn, S. G.…, & Hatfull, G. F. (2015). Whole genome comparison of a large collection of mycobacteriophages reveals a continuum of phage genetic diversity. Elife, 4, e06416.

Ramachandran, S., Kota, P., Ding, F., & Dokholyan, N. V. (2011). Automated minimization of steric clashes in protein structures. Proteins: Structure, Function, and Bioinformatics, 79(1), 261–270.

Rost, B., Yachdav, G., & Liu, J. (2004). The predictprotein server. Nucleic acids research, 32(Suppl_2), W321–W326.

Rule, C. S., Patrick, M., & Sandkvist, M. (2016). Measuring in vitro ATPase activity for enzymatic characterization. JoVE (Journal of Visualized Experiments), (114), e543.05.

Sambrook, J. (2000). Russell, D. ln Molecular Cloning: A Laboratory Manual.

Shen, H. B., & Chou, K. C. (2009). Predicting protein fold patterns with functional domain and sequential evolution information. Journal of Theoretical Biology, 256(3), 441 –446.

Sinha, A., Eniyan, K., Manohar, P., Ramesh, N., & Bajpai, U. (2020). Characterization and genome analysis of B1 sub-cluster mycobacteriophage PDRPxv. Virus Research, 279, 197884.

Skurnik, M., & Strauch, E. (2006). Phage therapy: facts and fiction. Intern ational Journal of Medical Microbiology, 296(1), 5 –14.

Snider, J., Thibault, G., & Houry, W. A. (2008). The AAA+ superfamily of functionally diverse proteins. Genome biology, 9(4), 216.

Stockley, P. G. (2009). Filter-binding assays. In DNA-Protein Interactions (pp. 1–14). Humana Press.

Thompson, J. D., Higgins, D. G., & Gibson, T. J. (1994). CLUSTAL W: improving the sensitivity of progressive multiple sequence alignment through sequence weighting, position-specific gap penalties and weight matrix choice. Nucleic acids research, 22(22), 4673–4680.

Vina, A. (2010). Improving the speed and accuracy of docking with a new scoring function, efficient optimization, and multithreading Trott, Oleg; Olson, Arthur J. J. Comput. Chem, 31(2), 455–461.

Yan, J., & Kurgan, L. (2017). DRNApred, fast sequence-based method that accurately predicts and discriminates DNA-and RNA-binding residues. Nucleic acids research, 45(10), e84 –e84.

Zhang, Y. (2008). I-TASSER server for protein 3D structure prediction. BMC bioinformatics, 9(1), 40.

